# Methyltransferase DNMT3B promotes colorectal cancer cell proliferation by inhibiting PLCG2

**DOI:** 10.1101/2024.05.13.594025

**Authors:** Yong Ji, Yang Wang, Jiacheng Zou, Guanghao Liu, Mingyu Xia, Jun Ren, Daorong Wang

**Affiliations:** The Yangzhou School of Clinical Medicine of Dalian Medical University, Yangzhou, 225001, China; Clinical Medical College, Yangzhou University, Yangzhou, 225001, China; Jingjiang People’s Hospital, Jingjiang, 214500, China; The Yangzhou Clinical Medical College of Xuzhou Medical University, Yangzhou, 225001, China; General Surgery Institute of Yangzhou, Yangzhou University, Yangzhou 225001, China, Yangzhou, 225001, China; Yangzhou Key Laboratory of Basic and Clinical Transformation of Digestive and Metabolic Diseases, Yangzhou, 225001, China

**Author notes:** Correspondence: Daorong Wang; Jun Ren. This article Ji Yong and Wang Yang have the same contribution.

**Keywords:** Epigenetics, DNA methylation, DNMT3B, PLCG2, colorectal cancer

## Abstract

**Background:** Aberrant DNA methylation patterns in the promoter region of PLCG2 have been associated with dysregulated signaling pathways and cellular functions. Its role in colorectal cancer cells is still unknown.

**Methods:** qRT-PCR was used to examine DNMT3B expression in colorectal cancer. Western blot and immunohistochemistry were used to analyze DNMT3B and PLCG2 protein levels in colorectal tissues and cell lines. The cell counting kit-8 and colony experiments were used to identify the proliferation of colorectal cancer cells. Methylation-specific PCR (MSP) and bisulfite-sequencing PCR (BSP) was used to measure DNA methylation levels.

**Results:** DNMT3B is overexpressed in colorectal cell in TCGA datasets and Kaplan-Meier plots. DNMT3B is significantly overexpressed in tumor tissues compared to adjacent non-tumor tissues. Western blotting results demonstrated high expression of DNMT3B in tumor tissues.

Compared to normal colonic epithelial cells, colorectal cancer cell lines exhibited elevated levels of PLCG2 methylation. oePLCG2 effectively prevented the in vivo xenograft tumor growth of colorectal cancer.

**Conclusions:** PLCG2 is identified as a key downstream regulatory protein of DNMT3B in colorectal cancer. DNMT3B Inhibits PLCG2 transcription through methylation of the PLCG2 promoter region. DNMT3B controls colorectal cancer cell proliferation through the PLCG2, which is useful for creating therapeutic approaches that target PLCG2 expression for the treatment of colorectal cancer.

## Introduction

Colorectal cancer (CRC) is the third most common and deadly tumor type worldwide, although great progress has been made in CRC treatment over the past few years[1]. While there have been some advancements in early diagnosis rates, the current surgical and chemoradiotherapy treatments are not optimal[2]. This is attributed to the pathological stage, invasion, and metastasis of colorectal cancer cells, which compromise the effectiveness of the treatment[3].

DNA methylation plays a crucial role in the initiation and progression of colorectal cancer[4–6]. It is an epigenetic modification involving the addition of methyl groups to DNA molecules. In colorectal cancer, DNA methylation typically involves the increased or decreased methylation of certain genes, potentially leading to changes in the expression levels of these genes. Generally, in colorectal cancer, tumor suppressor genes often experience increased methylation, suppressing their function and diminishing their ability to resist cancer[7]. Conversely, some oncogenes may undergo decreased methylation, resulting in their overactivation and promoting the growth and spread of cancer cells[8]. Furthermore, the patterns and extent of DNA methylation can serve as markers for the diagnosis and prognosis of colorectal cancer[9–11]. Therefore, gaining a deeper understanding of the role of DNA methylation in colorectal cancer not only helps unravel the molecular mechanisms of the disease but also provides crucial clues for the development of more effective treatment and diagnostic approaches.

DNMT3B (DNA Methyltransferase 3B) is a DNA methyltransferase belonging to the DNA methyltransferase family. This enzyme is responsible for the addition of methyl groups to DNA molecules, thereby influencing gene expression and cellular genomic stability. The abnormal activity of DNMT3B has been implicated in the occurrence and development of diverse cancers, specifically encompassing cervical cancer[12], breast cancer[13], endometrial cancer[14], and gastric cancer[15]. It induces abnormal DNA methylation, affecting the expression of tumor suppressor genes and oncogenes, consequently propelling tumor growth and advancement.

PLCG2, or Phospholipase C-γ2, is a protein belonging to the phospholipase C enzyme family. It serves as a substrate for BTK (Bruton’s tyrosine kinase), and the inactivation of BTK leads to phosphorylation of AKT[16]. The interplay between PLCG2 and AKT phosphorylation underscores the complexity of cellular signaling networks and provides insights into potential mechanisms of negative feedback regulation in cellular processes. Phospholipase C-γ2 (PLCG2), recognized as a pivotal signaling protein, plays a vital role in maintaining normal cellular function and immune responses[17]. In-depth investigation into PLCG2 is instrumental in enhancing our understanding of its functions within cellular biology and the mechanistic underpinnings of disease pathogenesis. This scrutiny not only contributes to unraveling the molecular intricacies of PLCG2 but also sheds light on its implications in cellular processes and disease mechanisms, providing valuable insights for potential therapeutic.

Understanding the molecular origins of colorectal cancer (CRC) is essential for enhancing the survival rates of CRC patients. Consequently, a more comprehensive understanding of the molecular irregularities linked to CRC pathogenesis has the potential to enhance treatment modalities and improve the overall survival of individuals affected by this condition.

## Materials and Methods

### Cell culture

The Cell Bank of the Chinese Academy of Sciences (Shanghai, China) provided the human colorectal cancer cell lines (HCT116, and SW620) and epithelial cell line NCM460, which were used in this investigation. The RPMI-1640 medium (HyClone, Logan, USA) used for cell culture was supplemented with 1% penicillin/streptomycin and 10% Fetal Bovine Serum (FBS) (Life Technologies, Waltham, USA). The cells were cultivated at 37°C in a humidified incubator with a 5% CO2 environment.

### Patient samples

From March 2021 to July 2022, eighty patients with each set of (cancer and normal) frozen colorectal cancer tissues were obtained from CRC excision at the Northern Jiangsu People’s Hospital. The patient’s CRC was confirmed at the time of their initial diagnosis, and neither had previously undergone radiotherapy or chemotherapy. The samples taken from the patients were placed in liquid nitrogen right away. The Medical Ethics Committee of Northern Jiangsu People’s Hospital provided its ethical approval (2020KY-137).

### Immunohistochemistry (IHC)

A tissue microarray (TMA) that included samples from 80 patients with histolog-ically confirmed colorectal cancer and 80 controls was generated according to a previ-ously described method. The sections were deparaffinized in xylene, rehydrated in a graded alcohol series and citrate buffer, and then blocked with 3% hydrogen perox-ide. Subsequently, the sections were incubated with a primary antibody directed against DNMT3B (1:100; 57868, Cell Signaling Technology, Massachusetts, USA) PLCG2 (1:150; 5690, Cell Signaling Technology, Massachusetts, USA) and then with a biotin-conjugated secondary antibody (SA1050; 7868 Boster, Wuhan, China), followed by incubation with a streptavidin–peroxidase complex. Five high-power fields (400× magnification) were selected randomly and photographed for each slide. The protein expression scoring was evaluated by taking both the proportion of positive cells [0 (<5%), 1 (5%–25%), 2 (26%–50%), 3 (51%–75%), and 4 (>75%)] and the intensity of cell staining [0 (negative), 1 (weak), 2 (moderate), and 3 (strong)] into account. The final staining scores were calculated by multiplying the staining intensity by the degree of staining. DNMT3B staining was considered low or high using a cutoff value of 5 based on the analysis from the receiver operating characteristic (ROC) curve. A final score greater than 5 was defined as a high expression of PLCG2.

### Quantitative real-time PCR (qRT-PCR)

Total RNA was extracted by utilizing TRIzol from Invitrogen (Carlsbad, USA), and a cDNA synthesis kit (K1622; Thermo Scientific, Waltham, USA) was used to reverse-transcribe 1 g of the extracted RNA. Each RT-PCR procedure required 200 ng of cDNA in total. Quantitative real-time PCR (qRT-PCR) was conducted through 2× Universal SYBR Green Fast qPCR Mix (ABclonal, Wuhan, China). The following amplification procedures were used: 95°C for 3 min, then 40 cycles of 95°C for 5 s and 60°C for 30s. The primer pair sequences are as follows: PLCG2 Forward Primer 5’-TCCACCACGGTCAATGTAGAT-3’, Reverse Primer 5’-CCCTGGGCGGATTTCTTTTAT-3’; DNMT3B Forward Primer 5’-AGGGAAGACTCGATCCTCGTC-3’, Reverse Primer 5’-GTGTGTAGCTTAGCAGACTGG-3’ and GAPDH Forward Primer 5’-ACGGATTTGGTCGTATTGGGCG-3’, Reverse Primer 5’-GCTCCTGGAAGATGGTGATGGG-3’. The 2−ΔΔCt method was employed to determine the target gene expression.

### Western blot analysis

Cells were lysed by utilizing RIPA buffer (Beyotime, Shanghai). Each specimen (30 mg of protein) was separated by SDS-PAGE just after protein concentration was determined. The separated proteins were then electro-transferred on 0.45-m polyvinylidene difluoride membranes (Millipore, Billerica, USA). The membranes were subsequently blocked using 5% skim milk for two hours at room temperature after being washed three times with TBST. A primary antibody was then incubated with the membranes at 4°C for an overnight period. The membranes were then incubated with a horseradish peroxidase-conjugated secondary antibody (1:5000; ABclonal, China) for two hours at room temperature after being washed three times with TBST. After being treated with a secondary antibody coupled to peroxidase, signals were discovered utilizing an enhanced chemiluminescence substrate (Milipore, Schwalbach, Germany). The bands on the Western blot were measured for intensity using Image J and for analysis using GraphPad Prisma 8. The primary antibodies used in this study were as follows: rabbit anti-DNMT3B (1:2000; 57868, Cell Signaling Technology, Massachusetts, USA), rabbit anti-PLCG2 (1:2000; 5690, Cell Signaling Technology, Massachusetts, USA), rabbit anti p-AKT (1:2000, 4060, Cell Signaling Technology, Massachusetts, USA); rabbit anti-CyclinD1 (1:2000, 2922, Cell Signaling Technology, Massachusetts, USA); mouse anti-GAPDH (1:5000, ABclonal, China)

### Cell transfection

We utilized lentivirus to induce the overexpression of PLCG2 in the HCT116 cell line, establishing stable colorectal cancer cell lines with elevated PLCG2 levels. Two modified lentiviral vectors were generated: Vector (lentivirus-EGFP-Puro) and PLCG2 (lentivirus-PLCG2-EGFP-Puro) from GeneChem (Shanghai, China). Subsequently, 1×10^4^ cells were seeded into each well of a 24-well plate 12 hours before viral infection. Lentivirus was introduced into each well for a 72-hour period, succeeded by puromycin screening to confirm the establishment of stable cell lines. Fluorescence microscopy was employed to detect the fluorescence signal in the cells. The process was performed in accordance with the outlined methodology.

### Cell Counting Kit -8 assay (Cell Proliferation)

The single-cell suspension was produced by digesting the transfected cells with 0.25% trypsin after two rounds of PBS washing after they had attained 80% confluency. Six parallel wells were created for the experiment, and the cells were injected into a 96-well plate at a density of 1500 cells per well in 100 μl of the medium. Each well received 10 μl of CCK-8 solution (CK13; Dojindo, Shanghai, China) after 24, 48, and 72 hours of incubation. The optical density (OD) value in each well was then determined using a microplate reader at a wavelength of 450 nm. Each experiment was conducted three times.

### Colony formation

To evaluate the ability of single cells to form colonies, 500 cells/well were added in 24-well plates in RPMI Medium 1640 basic with 10% FBS. Cells were incubated for 14 days at 37 °C. Afterward, cells were washed, fixed in 4% paraformaldehyde, and stained with 0.5% crystal violet in acetic acid 3% (v/ v) for 20 min at room temperature. Colonies were photographed and quantified under a light microscope.

### Methylation-specific PCR (MSP) and bisulfite-sequencing PCR (BSP)

Using online MetPrimer software (http://www.urogene.org/methprimer) to analysis of CpG islands and primer design of human PLCG2 promoters for methylation-specific PCR (MSP) and bisulfite-sequencing PCR (BSP). For MSP analysis of the human *PLCG2* primer, we used the methylated forward primer ATTTTCGAGTTTAGACGATTTTTTC and the reverse primer CGCAATCCAAATATTTACCGTA and the unmethylated forward primer TTTTGAGTTTAGATGATTTTTTTGT and the reverse primer ACCACAATCCAAATATTTACCATA. The PCR control for input DNA (Input) was performed with the forward primer CCTGGCGGGTAATTGTGAAGA and the reverse primer CTGCGTGCCAAAGAAGAAACT. After PCR amplification, the products were analyzed on a 2% agarose gel and observed under ultraviolet light, and densitometric analysis was performed using ImageJ software.

Bisulfite sequencing modification of mouse DDIT4 promoter by EpiArt DNA Methylation Bisulfite Kit (Vazyme Biotech, China). Genomic DNA of bisulfite - treated mouse skeletal muscle was amplified by PCR with the BSP forward primer TTAGGTTTTTTAAAGAGTTGGGATG and the reverse primer ACAAAAAAAATTCCCCAATATCATA. The BSP amplification product was purified by Vazyme Biotech (China) amplification product purification kit and cloned into pGEM-Tasy-vector system (Promega, USA). and the percentages of methylated cytosines over total cytosines were calculated.

### Xenotransplantation Experiment

A total of 10 4-week-old BALB/c male nude mice (weight, 18–22 g) were purchased from GemPharmatech Co. Ltd. (Nanjing, China) and raised in a pathogen-free laminar flow cabinet throughout the experiments under the following conditions: Controlled humidity (30–40%), a constant temperature of 25 ℃, a 12-h light/dark cycle and free access to food and water. The ethical approval (approval no. 202111022) to perform the animal experiments was obtained from the Ethics Committee for Animal Experiments of the Yangzhou University (Yangzhou, China). The experimental protocol was performed in accordance with the Laboratory Animal Guideline for Ethical Review of Animal Welfare. 4-week-old male BALB/c nude mice were randomly divided into Vector and oePLCG2 groups (n = 5 in both groups). Under isoflurane inhalation anesthesia (1–2%), ∼1 × 10^6^ HCT116 cells of stably transfected strains Vector/oePLCG2 resuspended in 100 µl PBS were subcutaneously into the left armpit of the mice. The health and behaviour of the mice were monitored every 2 days to determine if there were difficulties eating or drinking, unrelieved pain or distress without recovery. If the tumor reached 2000 mm3, the animal would be euthanized as a humane endpoint. The following formula was used to calculate the tumor volume (V) every week: V = (Width2 × Length)/2. Four weeks post-inoculation, all the mice were sacrificed by cervical dislocation under anesthesia. The method of anesthesia used for the mice was CO2 asphyxiation (CO2 was introduced into the chamber at a rate of 40–70% of the chamber volume per min to minimize distress). Dilated pupils were then used to verify death. Then, the tumors were removed and weighed.

### Statistical analysis

Statistical analysis was performed using the GraphPad Prism (Version 8.0). Quantitative data are presented as mean ± SD. Student’s t-test analysed differences in the mean of two samples. The association between DNMT3B expression and clinicopathological characteristics was analyzed by the Chi-square test (χ2). Patient survival was evaluated utilizing the Kaplan-Meier method. Counting and calculating area were executed with ImageJ (1.46r). All statistical tests were considered significant when \**P* <0.05, ** *P* <0.01, or *** *P* <0.001.

## Results

### DNMT3B is overexpressed in colorectal cell in TCGA datasets and Kaplan-Meier plots

We found that DNMT3B is significantly overexpressed in tumor tissues based on the TCGA database (https://www.cancer.gov/ccg/research/genome-sequencing/tcga/using-tcga-da ta/citing) (Fig 1A). Subsequently, we analyzed the effect of DNMT3B on survival using the Kaplan-Meier plotter database (kmplot.com). The results showed that patients with high expression of DNMT3B exhibited poorer prognosis (see Fig 1B). To validate the expression level of DNMT3B in clinical tumor tissues, we collected clinical tissue samples from 80 patients with a history of colorectal cancer surgery at Subei People’s Hospital. Tissue microarrays were constructed, and immunohistochemical staining was performed. The results showed that DNMT3B was significantly overexpressed in tumor tissues compared to adjacent non-tumor tissues (see Fig 1C). Additionally, fresh tissue samples were collected for protein extraction, and Western blotting results demonstrated high expression of DNMT3B in tumor tissues (see Fig 1D, E).

**Fig 1.**
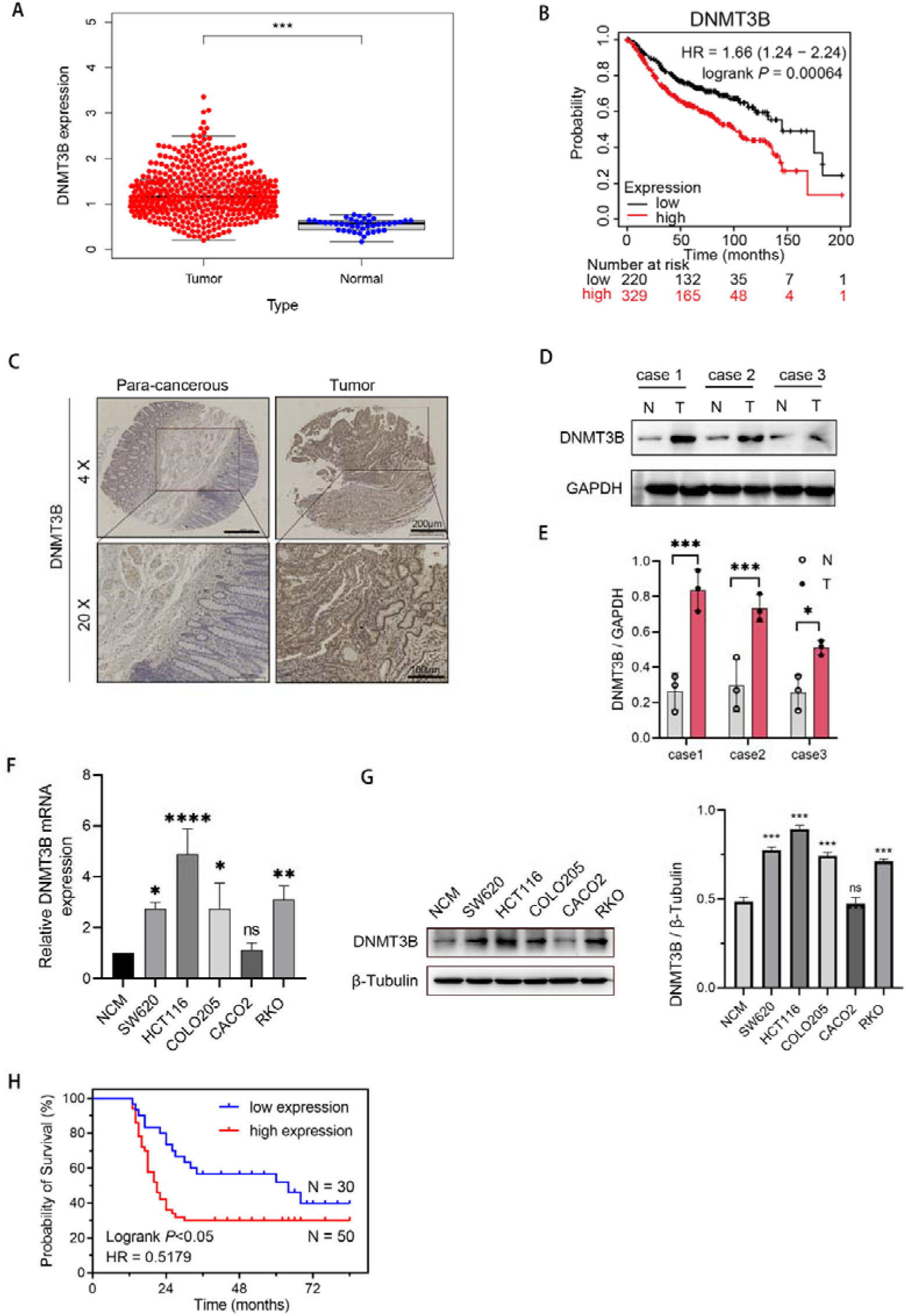
A The expression of DNMT3B in colorectal cancer was analyzed by TCGA database B Kaplan-Meier plotter database to analyze the effect of DNMT3B on prognosis in colorectal cancer C Immunohistochemical analysis of DNMT3B in paired noncancerous tissues and CRC tissues of patients D The protein expression level of DNMT3B in paired noncancerous tissues and CRC tissues of patients E Quantitative analysis of western blot F The mRNA expression level of DNMT3B in qRT-PCR colorectal cancer cells and epithelial cell line NCM460 G The protein expression level of DNMT3B in western blot colorectal cancer cells and epithelial cell line NCM460 H Kaplan-Meier survival analysis shown DNMT3B overexpression predicted poorer prognosis in colorectal cancer.

To further investigate the role of DNMT3B in tumor progression, we examined the expression level of DNMT3B in common colorectal cancer cell lines. The results revealed that compared to the colon epithelial cell line NCM460, the mRNA expression of DNMT3B was elevated in colorectal cancer cell lines SW620, HCT116, COLO205, and RKO (see Fig 1F). DNMT3B exhibited a similar trend at the protein level (see Fig 1G). Our follow-up data suggest that patients with high expression of DNMT3B exhibited a shorter survival duration compared to those with low expression of DNMT3B (see Fig 1H).

### DNMT3B is highly expressed in colorectal cancer

DNMT3B positivity staining is challenging to detect in normal tissues but readily observable in colorectal cancer tissues. Utilizing ROC analysis, an IHC score of 5 serves as the cutoff point to distinguish between high and low expression levels of DNMT3B. The high expression rate of DNMT3B in colorectal cancer was 80%, which was significantly higher than that in normal tissue(Table 1). We examined the correlation between DNMT3B expression and clinical-pathological characteristics of colorectal cancer. High expression of DNMT3B significantly correlates with T staging, lymph node metastasis, tumor size, and distant metastasis, while showing no significant association with age, BMI, or gender (Table 2).

**Table 1.** DNMT3B expression was detected in colorectal cancer by IHC.

| Types | N | DNMT3B |  | P-value |
| --- | --- | --- | --- | --- |
|  |  | Low expression | High expression |  |
| Tumor tissues | 40 | 8 (20.0%) | 32 (80.0%) | 0.001 |
| Normal tissues | 40 | 22 (55.0%) | 18 (45.0%) |  |

**Table 2.** The correlation between DNMT3B expression and clinicopathological characters was analyzed in colorectal cancer.

| Types | Low expression<br>(n=30) | High expression<br>(n=50) | P-value |
| --- | --- | --- | --- |
| **Gender** |  |  | 0.441 |
| Male [n(%)] | 10(12.5%) | 21(26.3%) |  |
| Female [n (%)] | 20(25.0%) | 29(36.3%) |  |
| **Age [n (SD)]** | 59.97(13.4) | 58.92(10.8) | 0.702 |
| **BMI (kg/m<sup>2</sup>) [n (SD)]** | 23.9(2.7) | 24.5(3.5) | 0.448 |
| **T [n (%)]** |  |  | 0.026 |
| 1 | 2(2.5%) | 4(5.0%) |  |
| 2 | 15(18.8%) | 21(26.3%) |  |
| 3 | 13(16.3%) | 13(16.3%) |  |
| 4 | 0 | 12(15.0%) |  |
| **N [n (%)]** |  |  | 0.006 |
| 0 | 16(20.0%) | 10(12.5%) |  |
| 1 | 11(13.8%) | 26(32.5%) |  |
| 2 | 3(3.8%) | 14(17.5%) |  |
| **M [n (%)]** |  |  | 0.044 |
| 0 | 25(31.3%) | 31(38.8%) |  |
| 1 | 5(6.3%) | 19(23.8%) |  |
| **TNM [n (%)]** |  |  | 0.018 |
| 1 | 12(15.0%) | 8(10.0%) |  |
| 2 | 4(5.0%) | 2(2.5%) |  |
| 3 | 9(11.3%) | 21(26.3%) |  |
| 4 | 5(6.3%) | 19(23.8%) |  |
| **Tumor size (cm) [n (SD)]** | 3.2(1.0) | 4.6(1.4) | 0.000 |

### PLCG2 is identified as a key downstream regulatory protein of DNMT3B in colorectal cancer

To better delineate the downstream regulatory proteins of DNMT3B in colorectal cancer, we treated HCT1116 colon cancer cells with the DNMT3B inhibitor SGI-1027 (SGI) for 24 hours, collected the cells, and performed transcriptomic sequencing. Principal component analysis was conducted on the differentially expressed genes after SGI treatment, revealing distinct clustering of the SGI-treated group (see Fig 2A). Heatmap analysis of the differentially expressed genes showed significant upregulation of PLCG2 protein and notable differences in the PI3K-AKT signaling pathway (see Fig 2B). Subsequently, we performed KEGG pathway enrichment analysis on the differentially expressed genes, which indicated a significant impact on the tumor cell cycle (see Fig 2C). To investigate the clinical relevance of PLCG2, we conducted immunohistochemical staining of PLCG2 using tissue microarrays. The results showed stable expression of PLCG2 in normal tissues, whereas PLCG2 protein expression was decreased in colorectal cancer tissues (see Fig 2D). Similarly, we validated the expression of PLCG2 in TCGA, revealing a significant downregulation of PLCG2 expression in colorectal cancer tumor tissues (see Fig 2E).

**Fig 2.**
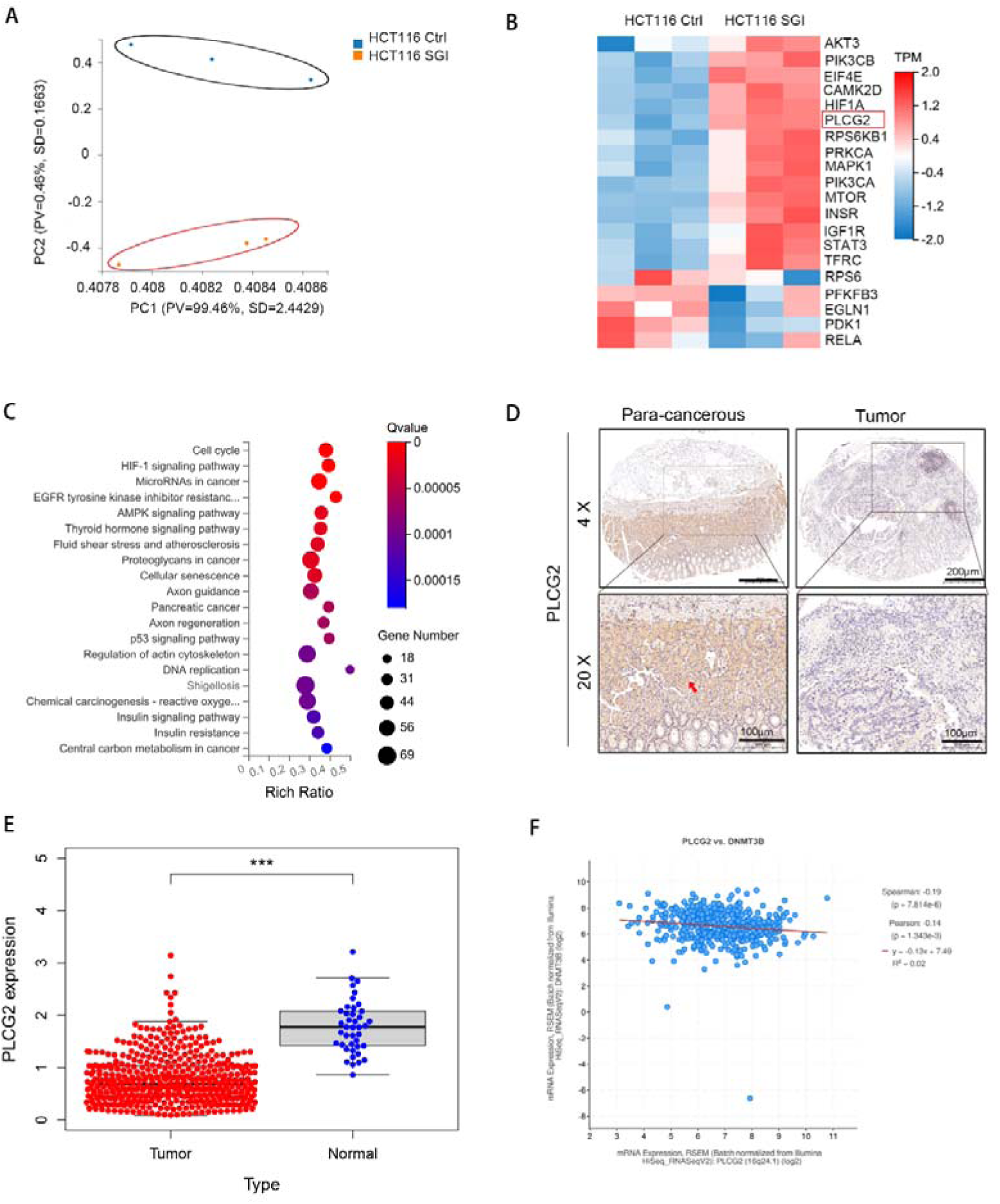
A Principal component analysis was conducted on the differentially expressed genes after SGI treatment for 24 hours B Heat map analysis of HCT116 cells after SGI treatment for 24 hours C KEGG pathway enrichment analysis on the differentially expressed genes D Immunohistochemical analysis of PLCG2 in paired noncancerous tissues and CRC tissues of patients E The expression of PLCG2 in colorectal cancer was analyzed by TCGA database F We analyzed the correlation between DNMT3B and PLCG2 expression in the TCGA database

Furthermore, we analyzed the correlation between DNMT3B and PLCG2 expression in the TCGA database, revealing a significant negative correlation between DNMT3B and PLCG2 (see Fig 2F). These results suggest that DNMT3B may exert oncogenic effects by downregulating PLCG2.

### PLCG2 overexpression inhibits tumor cell proliferation in Colorectal cells

To determine the role of PLCG2 in colorectal cancer, we constructed a lentiviral vector overexpressing PLCG2 and obtained stable overexpression cell lines. Plate colony formation assays were conducted on cells overexpressing PLCG2, revealing a reduction in tumor cell proliferation upon overexpression of PLCG2 protein (see Fig 3A). CCK8 assays were performed on cells overexpressing PLCG2, showing decreased proliferation at 48, 72, and 96 hours compared to the control group (see Fig 3B).

**Fig 3.**
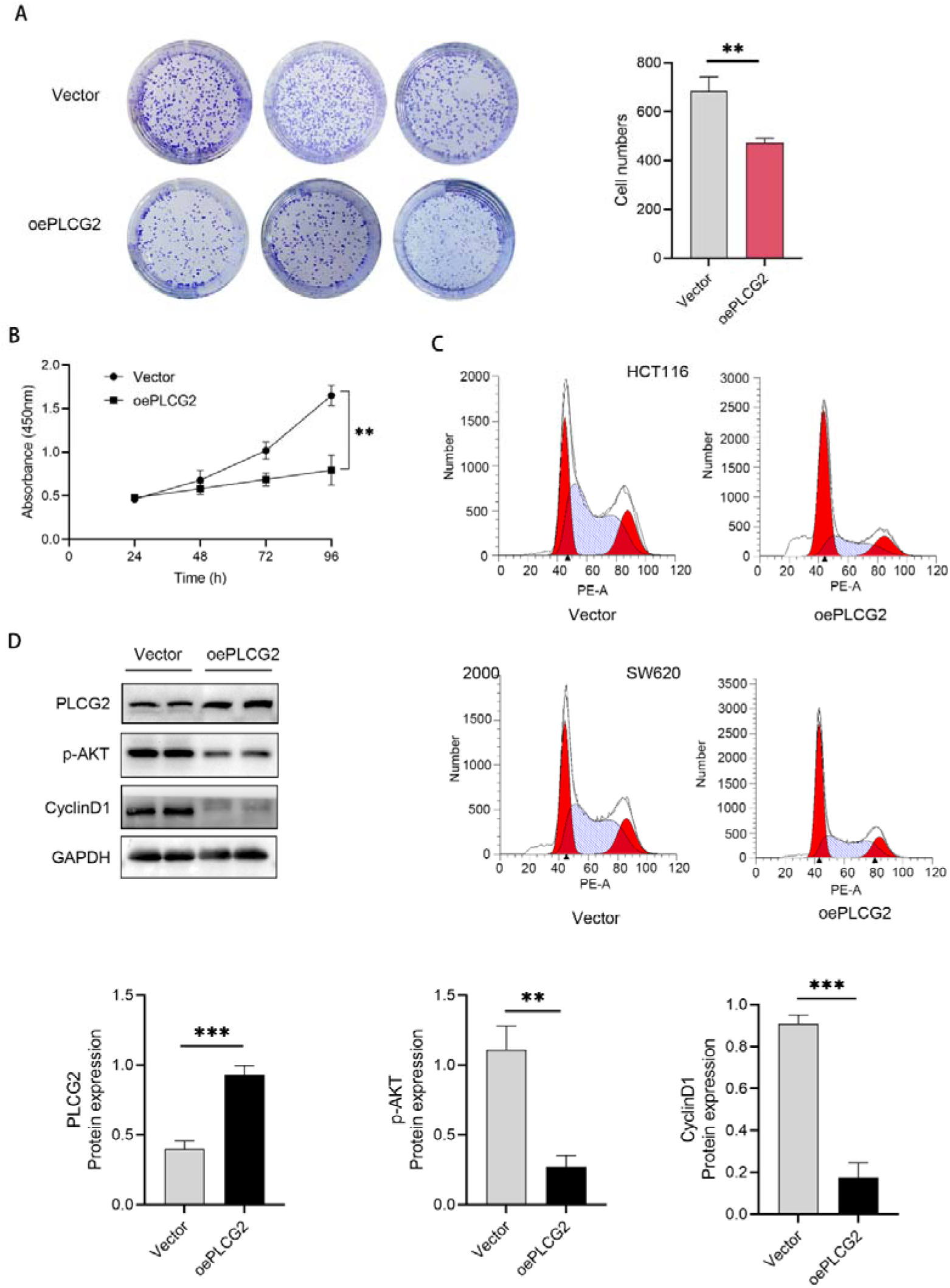
A Detection of the proliferation of HCT116 cells after PLCG2 overexpression by colony formation assay B The proliferation of vector cells and oePLCG2 cells in HCT116 was determined by CCK-8 assay C The cell cycle of vector cells and oePLCG2 cells in HCT116 and SW620 was detected by flow cytometry D The protein level was detected by western blot in HCT116 cells

Subsequently, flow cytometry cell cycle analysis was conducted on SW620 and HCT116 cells overexpressing PLCG2, revealing a significant increase in the proportion of cells in the G1 phase after overexpression of PLCG2 (see Fig 3C). Protein was extracted from HCT116 cells overexpressing PLCG2 and subjected to Western blotting analysis, which showed a significant decrease in the expression levels of downstream proteins p-AKT and Cyclin D1 after overexpressing PLCG2 (see Fig 3D).

### DNMT3B Inhibits PLCG2 transcription through methylation of the PLCG2 promoter region

To elucidate the molecular mechanism by which DNMT3B regulates PLCG2, we analyzed the methylation sites in the PLCG2 promoter region using the Methyprimor database. Unexpectedly, a significant CpG island was found 1000bp upstream of the PLCG2 transcription start site (Fig 4A), indicating that DNMT3B may suppress PLCG2 expression through DNA methylation. Methylation-specific PCR amplification was conducted using methy and unmet primers designed for the PLCG2 promoter region after bisulfite modification. Compared to normal colonic epithelial cells, colorectal cancer cell lines exhibited elevated levels of PLCG2 methylation and decreased levels of non-methylation (Fig 4B). To investigate the role of DNMT3B in methylation, cells were treated with the inhibitor SGI for 24 hours, genomic DNA was extracted, and MSP experiments were performed. The results showed decreased methylation levels in colorectal cancer cells treated with SGI (Fig 4C). Similarly, cell proteins were extracted using the same treatment method, and Western blot analysis revealed that PLCG2 protein levels increased after treatment with the DNMT3B inhibitor SGI (Fig 4E).

**Fig 4.**
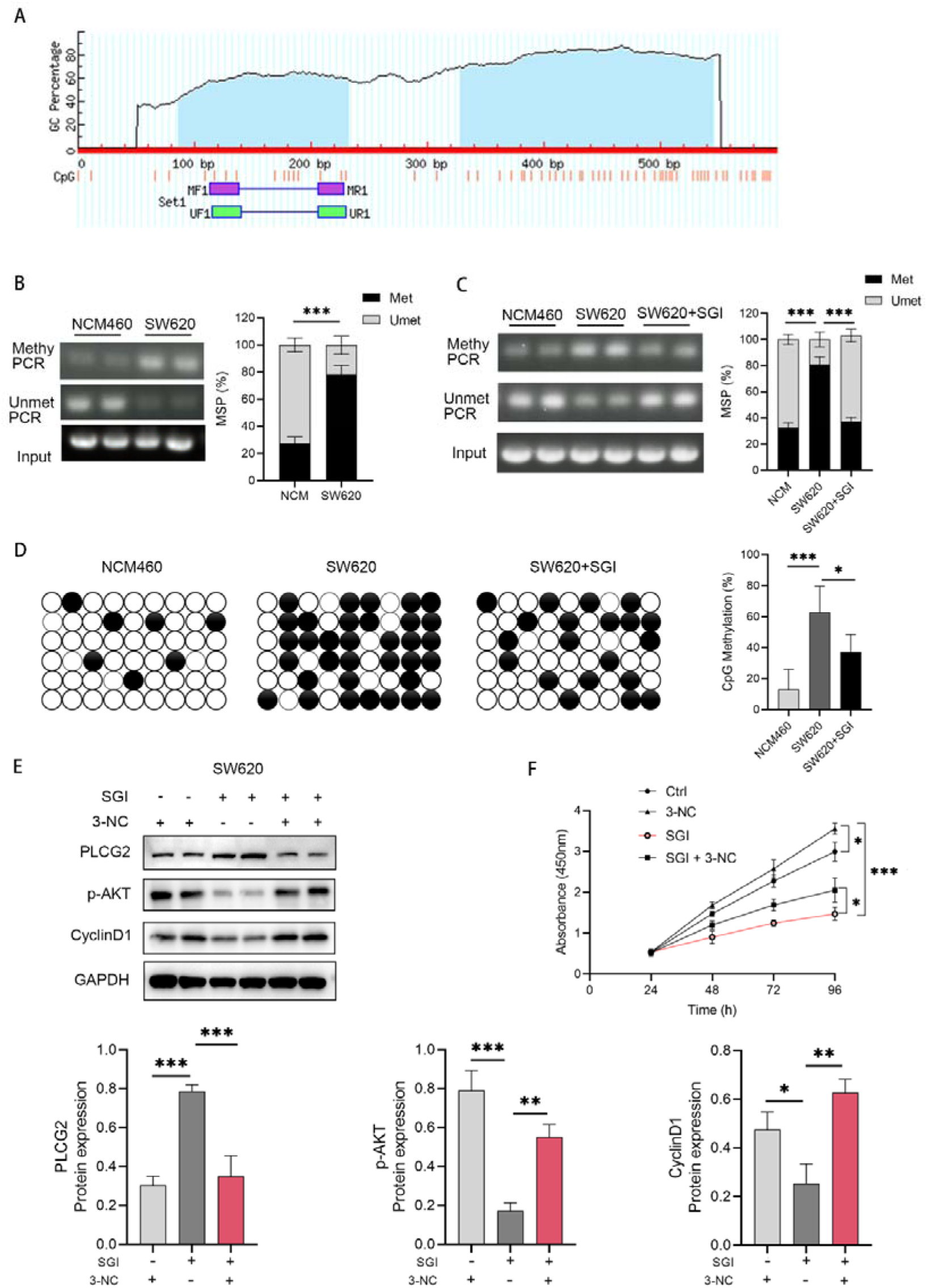
A We analyzed the methylation sites in the PLCG2 promoter region using the Methyprimor database B Methylation-specific PCR amplification was conducted using methy and unmet primers designed for the PLCG2 promoter region after bisulfite modification. C MSP analysis of SW620 cells were treated with the SGI for 24 hours D BSP analysis of SW620 cells were treated with the SGI for 24 hours E In rescue experiments, PLCG2 protein levels increased after SGI treatment, while the use of 3-NC reduced PLCG2 activity, resulting in the restoration of downstream p-AKT and CyclinD1 expression F The same methods were applied in CCK8 experiments, where cell viability significantly decreased after 24 hours of SGI treatment. Moreover, the proliferation capacity of cells was higher in the group treated with both SGI and 3-NC compared to cells treated solely with SGI

PLCG2 is a substrate of Bruton’s tyrosine kinase (BTK), and decreased BTK levels result in AKT phosphorylation. Elevated PLCG2 levels lead to decreased p-AKT levels and reduced CyclinD1 levels, causing cell cycle arrest at the G1 phase (Fig 4E). 3-NC is an inhibitor of PLCG2. In rescue experiments, PLCG2 protein levels increased after SGI treatment, while the use of 3-NC reduced PLCG2 activity, resulting in the restoration of downstream p-AKT and CyclinD1 expression (Fig 4E). The same methods were applied in CCK8 experiments, where cell viability significantly decreased after 24 hours of SGI treatment. Moreover, the proliferation capacity of cells was higher in the group treated with both SGI and 3-NC compared to cells treated solely with SGI (Fig 4F).

### Overexpression of PLCG2 was found to inhibit the formation of colorectal cancer tumors in a xenograft model

Immunodeficient BALB/c mice harboring HCT116 cells, which were stably transfected with either the vector or oePLCG2 lentivirus, were employed to investigate the role of PLCG2 in colorectal cancer carcinogenesis in vivo. This was done to further validate whether PLCG2 functions in in vivo models. Five-week-old nude mice were subcutaneously injected with HCT116 vector cells and HCT116 oePLCG2 cells. A palpable mass emerged at the injection site one week post-injection. After 4 weeks, the tumors were harvested and examined (Fig 5A). The largest and smallest diameters of the masses were measured weekly. As expected, starting from the third week, the overexpression of PLCG2 significantly inhibited the growth of HCT116 tumors in mice compared to the control group (Fig 5B). After the fourth week, the nude mice were euthanized, and tumor weights and volumes were assessed. The average weight of the oePLCG2 group (255.80 ± 21.49 mg) was significantly lower than that of the vector group (555.80 ± 27.70 mg) (Fig 5C, P < 0.001). Moreover, the average volume of the oePLCG2 group was markedly smaller than that of the vector group (Fig 5D, P < 0.001) (279.20 ± 18.59 vs. 580.80 ± 12.25 mm3). The expression of protein in mouse tumor tissue was analyzed by western blot. The results showed that the level of p-AKT CyclinD1 protein was significantly reduced after PLCG2 overexpression in subcutaneous tumor tissues of nude mice (Fig 5E). The findings demonstrated that oePLCG2 effectively prevented the in vivo xenograft tumor growth of colorectal cancer.

**Fig 5.**
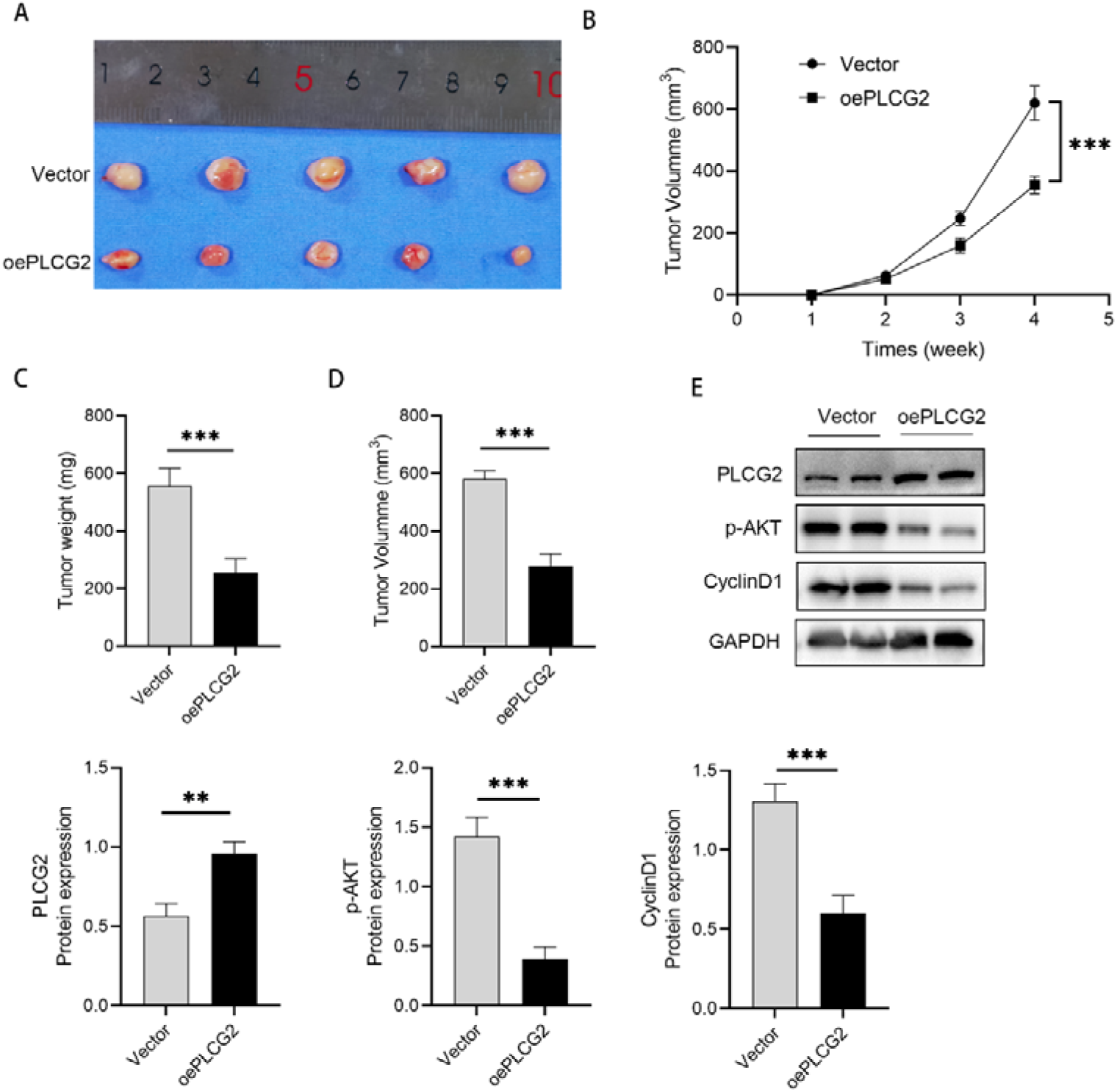
Effects of oePLCG2 on tumorigenesis in nude mice in vivo. A Xenograft models in nude mice were generated using HCT116 cells transfected with Vector (n=5) or oePLCG2 (n=5). B Subcutaneous tumor growth curve in nude mice. C Detection of the tumor weight of HCT116 cells transfected with PLCG2 lentiviral OE vector after euthanasia. D Detection of the tumor volume of HCT116 cells transfected with PLCG2 lentiviral OE vector after euthanasia. E The expression of protein in mouse tumor tissue was analyzed by western blot.

## Discussion

Colorectal cancer poses a significant threat characterized by a grim prognosis, delayed diagnosis, and insufficient treatment[1]. Early detection of colorectal cancer is infrequent due to vague symptoms that often manifest later in the clinical progression. The pathogenesis of colorectal cancer is intricate. For instance, aberrant regulation of transcription factors in colorectal cancer may result in abnormal gene expression, impacting processes such as cell proliferation, differentiation, and apoptosis[18]. Additionally, the dysfunction of mitochondria and alterations in cellular energy metabolism may play a role in the onset of colorectal cance[19]. Ubiquitination is a mechanism associated with protein degradation, and excessive ubiquitination could lead to abnormal degradation of crucial regulatory proteins, thereby influencing cellular functions[20]. The precise pathogenic mechanism of colorectal cancer remains incompletely elucidated. The mechanism involves the interplay of multiple factors, and ongoing research is continually advancing to provide a more comprehensive understanding of the disease formation process. Consequently, the identification of novel molecular pathways and potential therapeutic targets becomes imperative to mitigate the severity of colorectal cancer.

In this research, we uncovered several significant discoveries that contribute to a deeper comprehension of the epigenetic mechanisms involved in colorectal cancer. The investigation into the relationship between DNMT3B and PLCG2 represents a crucial exploration in the field of molecular biology. DNMT3B, as a vital member of the DNA methyltransferase family, is responsible for adding methyl groups to DNA molecules, influencing gene expression, and maintaining genomic stability[21]. On the other hand, PLCG2 is a key protein involved in cellular signaling pathways[22–24]. Recent research has sparked an in-depth examination of the potential interactions and cross-pathways between DNMT3B and PLCG2, aiming to elucidate their mutual impact on cellular processes and functions. Understanding the relationship between these two molecules becomes paramount, as it may clarify the intricate mechanisms governing gene regulation, epigenetic modifications, and signal transduction pathways. This study aims to comprehensively overview the current research status regarding the relationship between DNMT3B and PLCG2. The exploration encompasses a detailed scrutiny of their individual functionalities, the molecular mechanisms underlying their interactions, and the biological consequences of their mutual influence.

The clinical relevance of DNMT3B in colorectal cancer has been a focus of research, with efforts directed towards assessing its potential as a diagnostic biomarker or therapeutic target[25–27]. The development of anti-tumor agents has consistently been a central theme in clinical research[28]. In this domain, the focus of these endeavors is directed towards promoting the development of innovative drugs, with the aim of providing more effective and targeted therapeutic options for cancer patients. The exploration of DNMT3B inhibitors and their efficacy in mitigating colorectal cancer progression further signifies the translational implications of this research.

DNA methylation in the promoter region of PLCG2 is known to influence its expression levels. Aberrant methylation patterns in this gene have been associated with dysregulated signaling pathways and cellular functions in various cancers, including colorectal cancer. Alterations in DNA methylation within the regulatory regions of PLCG2 may lead to changes in its expression levels. This, in turn, can affect downstream signaling cascades and contribute to the progression of colorectal cancer. Association with Tumor Suppression: Studies suggest that DNA methylation-mediated silencing of PLCG2 may interfere with its tumor-suppressive functions. Understanding the specific CpG sites involved and the extent of methylation alterations is crucial for deciphering its role in colorectal carcinogenesis[29–31].

DNA methylation patterns in PLCG2 may serve as potential diagnostic biomarkers for colorectal cancer. Analyzing these epigenetic modifications could aid in early detection and risk stratification. The correlation between DNA methylation status in PLCG2 and clinical outcomes could establish its utility as a prognostic indicator. Stratifying patients based on these epigenetic profiles may guide personalized treatment approaches.

As research on DNA methylation in PLCG2 advances, further investigations are warranted to unravel the complexities of its epigenetic regulation, interactions with other signaling pathways, and functional consequences in colorectal cancer. The integration of these findings into clinical practice holds the potential to enhance diagnostic accuracy, prognostic precision, and the development of targeted epigenetic therapies for colorectal cancer patients.

## Conclusion

PLCG2 is identified as a key downstream regulatory protein of DNMT3B in colorectal cancer. DNMT3B Inhibits PLCG2 transcription through methylation of the PLCG2 promoter region. DNMT3B controls colorectal cancer cell proliferation through the PLCG2, which is useful for creating therapeutic approaches that target PLCG2 expression for the treatment of colorectal cancer.

## Acknowledgments

This work was supported by a fund was supported by Key Laboratory of Basic and Clinical Transformation of Digestive and Metabolic Diseases, Yangzhou, China (YZ2020159) and Application of Fluorescence Imaging in Minimally Invasive Surgery for Low Rectal Cancer (ZD2022047).

## Funding

The National Natural Science Foundation of China (No. 81972269) and Key Laboratory of Basic and Clinical Transformation of Digestive and Metabolic Diseases, Yangzhou, China (YZ2020159) and Application of Fluorescence Imaging in Minimally Invasive Surgery for Low Rectal Cancer (ZD2022047).

## Data Availability Statement

The data presented in this study are available on reasonable request.

## Competing interests

The authors declare that they have no competing interests.

## Medical Ethics

It’s approved by the ethics committee of Northern Jiangsu People’s Hospital, Yangzhou, China.

## Contribution

Yong Ji and Yang Wang: Conceptualization, study design, study proposal, manuscript writing, Figure legends and submission.

Jiacheng Zou & Guanghao Liu: Data extraction and Data analysis. Mingyu Xia: Manuscript revision and formatting.

Jun Ren & Daorong Wang: Study guidance and final review.

## Notes

### Competing Interest Statement

The authors have declared no competing interest.

